# Human-specific genomic features of pluripotency regulatory networks link NANOG with fetal and adult brain development

**DOI:** 10.1101/022913

**Authors:** Gennadi V. Glinsky

## Abstract

Genome-wide proximity placement analysis of diverse families of human-specific genomic regulatory loci (HSGRL) identified topologically-associating domains (TADs) that are significantly enriched for HSGRL and termed rapidly-evolving in humans TADs (revTADs; Genome Biol Evol. 2016 8; 2774-88). Here, human-specific genomic features of pluripotency regulatory networks in hESC have been analyzed. The primary focus was on identification of human-specific elements of the interphase chromatin architecture of TADs responsible for transcriptional regulatory control of the NANOG, POU5F1, and POU3F2 genes. Comparative analyses of the four adjacent TADs spanning ~3.3 Mb NANOG locus-associated genomic region were carried-out to highlight primate-specific genomic features. Lastly, the putative mechanisms of the genome-wide regulatory effects of human-specific NANOG-binding sites (HSNBS) on expression of genes implicated in the fetal and adult brain development have been examined. Acquisition of primate-specific regulatory loci appears to rewire TADs exerting transcriptional control on pluripotency regulators, revealing a genomic placement pattern consistent with the enhanced regulatory impact of NANOG in primates. Proximity placement analysis of HSNBS identified a large expression signature in the human fetal neocortex temporal lobe comprising 4,957 genes, which appear to retain acquired in the embryo expression changes for many years of human brain development and maintain highly concordant expression profiles in the neocortex and prefrontal cortex regions of adult human brain. Collectively, reported herein observations indicate that genomic elements of pluripotency regulatory circuitry associated with HSNBS, specifically proteins of the classical NurD chromatin remodeling complex, contribute to transcriptional regulation of a large set of genes implicated in development and function of human brain.

**List of abbreviations:** 5hmC, 5-Hydromethylcytosine

CTCF, CCCTC-binding factor

DHS, DNase hypersensitivity sites

FHSRR, fixed human-specific regulatory regions

GRNs, genomic regulatory networks

HAR, human accelerated regions

hCONDEL, human-specific conserved deletions

hESC, human embryonic stem cells

HSGRL, human-specific genomic regulatory loci

HSNBS, human-specific NANOG-binding sites

HSTFBS, human-specific transcription factor-binding sites

LAD, lamina-associated domain

LINE, long interspersed nuclear element

lncRNA, long non-coding RNA

LTR, long terminal repeat

MADE, methylation-associated DNA editing

mC, methylcytosine

mESC, mouse embryonic stem cells

NANOG, Nanog homeobox

nt, nucleotide

POU5F1, POU class 5 homeobox 1

PSDS, partial strand displacement state

TAD, topologically associating domains

TE, transposable elements

TF, transcription factor

TSC, triple-stranded complex

TSS, transcription start sites

SE, super-enhancers

SED, super-enhancer domains

sncRNA, small non coding RNA

## Introduction

The identity of embryonic stem cells (ESC) critically depends on continuing actions of key master transcription factors (Young, 2011; Ng and Surani, 2011), which govern the maintenance of the ESC’s pluripotency state by forming constitutively active super-enhancers (Hnisz et al., 2013; Whyte et al., 2013; Dowen et al., 2014). The most recent experiments show that regulation of pluripotency in ESC occurs within selected topologically-associating domains (TADs) by establishing specific sub-TAD structures, which are designed to isolate super-enhancers and target genes within insulated genomic neighborhoods (Dowen et al., 2014). These sub-TAD structures designated super-enhancer domains (SEDs) are formed by the looping interactions between two CTCF-binding sites co-occupied by cohesin. In ESC, segregation of super-enhancers (SEs) and target genes within TADs and SEDs is designed to isolate critical elements of the core pluripotency regulatory circuitry within insulated genomic neighborhoods to facilitate the coordination and enhance the precision of regulatory interactions between SE, SE-driven cell identity genes and repressed genes encoding lineage-specifying developmental regulators (Dowen et al., 2014). The sustained expression of key cell identity genes and repression of genes encoding lineage-specifying developmental regulators is essential for maintaining ESC identity and pluripotency state. These processes are governed by the master transcription factors (TFs), namely OCT4 (POU5F1), SOX2, and NANOG (OSN), that function by establishing SEs regulating cell identity genes, including master TFs themselves (Hnisz et al., 2013; Whyte et al., 2013).

Conventional enhancers comprise discrete DNA segments occupying a few hundred base pairs of the linear DNA sequence and harboring multiple transcription facor binding sites (TFBS). SEs consist of clusters of conventional enhancers that are densely occupied by the master transcription factors and Mediator (Whyte et al., 2013). It has been suggested that creation of new TFBS and increasing density of TFBS would increase the probability of the emergence of new enhancer elements during evolution (Glinsky, 2015-2017). In turn, creation of new enhancers and increasing their density would facilitate the emergence of new SE and SED structures. This sequence of events would imply that evolutionary time periods required for creation of TFBS, enhancers, and SEs are shortest for TFBS, intermediate for enhancers, and longest for SEs. Consistent with the model predictions, the estimated creation time is markedly longer for SEs compared to enhancers for conserved (8-fold), primate-specific (17-fold), and human-specific (63-fold) regulatory sequences (Glinsky, 2015-2017). Furthermore, the estimated creation time periods for enhancers appear several folds longer compared to the creation time estimates for TFBS sequences in both chimpanzee and humans (Glinsky, 2015-2017). Sequence conservation analyses demonstrated that 87% of new SEs and 75% of new enhancers in the hESC genome were created within evolutionary conserved sequences (Glinsky, 2017). Similarly, a majority (67%) of new TFBS for NANOG (59,221 of 88,351 binding sites) and CTCF (58,865 of 87,883 binding sites) proteins detected in the hESC genome compared with mouse ESC are located within conserved sequences (Glinsky, 2015-2017). Collectively, these observations are highly consistent with the idea that primate-specific genomic regulatory elements have rewired the core pluripotency regulatory networks of hESC (Kunarso et al., 2010).

Genome-wide meta-analysis of TFBS in hESC demonstrated that transposable element-derived sequences, most notably LTR7/HERVH, LTR5_Hs, and L1HS, harbor thousands of candidate human-specific transcription factor-binding sites, HSTFBS (Glinsky, 2015). A majority of these candidate human-specific regulatory loci (HSGRL) appears to function as TFBS for NANOG, POU5F1 (OCT4), and CTCF proteins exclusively in hESC, suggesting their critical regulatory role during the early-stage embryogenesis (Glinsky, 2015). Bioinformatics and proximity placement analyses revealed that hESC-specific NANOG-binding sites are enriched near the protein-coding genes regulating brain size, pluripotency long non-coding RNAs, hESC enhancers, and 5-hydroxymethylcytosine-harboring sequences immediately adjacent to TFBS. Candidate human-specific TFBS are placed near the coding genes associated with physiological development and functions of nervous and cardiovascular systems, embryonic development, behavior, as well as development of a diverse spectrum of pathological conditions such as cancer, diseases of cardiovascular and reproductive systems, metabolic diseases, multiple neurological and psychological disorders (Glinsky, 2015). These observations indicate that HSTFBS for NANOG, POU5F1, and CTCF proteins may represent functionally-significant human-specific genomic regulatory loci of the core pluripotency networks in hESC.

Despite recent significant progress, the precise genetic and molecular basis defining unique to human phenotypic features remains largely unknown and the hypothesis that unique to human phenotypes result from human-specific changes to genomic regulatory sequences (King and Wilson, 1975) remains highly relevant. The near-comprehensive catalogue of changes within protein-coding genes that occurred during human evolution suggested that these changes alone cannot explain all unique to human traits (Chimpanzee Sequencing and Analysis Consortium, 2005; Green et al., 2010; Meyer et al., 2012; Prüfer et al., 2012; 2014; Fu et al., 2014), supporting the idea that the key regulatory DNA sequences responsible for development of human-specific phenotypes are most likely located within the non-protein-coding genomic regions.

Extensive search for human-specific genomic regulatory loci (HSGRL) revealed thousands candidate HSGRL, a vast majority of which is residing within non-protein coding genomic regions (McLean et al., 2011; Shulha et al., 2012; Konopka et al., 2012; Capra et al., 2013; Marnetto et al., 2014; Glinsky, 2015-2017; Dong et al., 2016). Candidate HSGRL comprise multiple distinct families of genomic regulatory elements, including regions of human-specific loss of conserved regulatory DNA termed hCONDEL (McLean et al., 2011); human-specific epigenetic regulatory marks consisting of H3K4me3 histone methylation signatures at transcription start sites in prefrontal neurons (Shulha et al., 2012); human-specific transcriptional genetic networks in the frontal lobe (Konopka et al., 2012); conserved in humans novel regulatory DNA sequences designated human accelerated regions, HARs (Capra et al., 2013); fixed human-specific regulatory regions, FHSRR (Marnetto et al., 2014); DNase I hypersensitive sites (DHSs) that are conserved in non-human primates but accelerated in the human lineage (haDHS; Gittelman et al. 2015); human-specific transcription factor-binding sites, HSTFBS (Glinsky, 2015), and DHSs under accelerated evolution, ace-DHSs (Dong et al., 2016). Several experimentally testable hypotheses on biogenesis of HSGRL have been proposed and initial evidence of putative human-specific biological functions during development and pathological conditions for several HSGRL families have been reported (Pollard et al. 2006; Prabhakar et al. 2006; 2008; Bird et al. 2007; Capra et al., 2013; Marnetto et al., 2014; Glinsky, 2015-2017; Dong et al., 2016).

Using DNA sequences of the reference genome databases for multiple evolutionary diverse species, a baseline of the evolutionary expected rate of base substitutions can be estimated based on the experimentally determined level of conservation between multiple species at the given locus. This locus-specific baseline substitution rate then can be compared to the observed substitution rates on a lineage of interest in relation to the evolutionary expected rate of substitutions and the statistical significance of the difference can be estimated. This method appears particularly effective for identifying highly conserved sequences within noncoding genomic regions that have experienced a marked increase of substitution rates on a particular lineage. It has been successfully applied to humans (Pollard et al. 2006; Prabhakar et al. 2006; Bird et al. 2007), where the rapidly-evolving sequences that are highly conserved across mammals and have acquired many sequence changes in humans since divergence from chimpanzees were designated as human accelerated regions (HAR). Experiments attempting to interrogate HAR’s bioactivity revealed that some HARs function as non-coding RNA genes expressed during the neocortex development (Pollard et al. 2006) and human-specific developmental enhancers (Prabhakar et al. 2008). Recent computational analyses and transgenic mouse experiments demonstrated that many HARs represent developmental enhancers (Capra et al., 2013), supporting the hypothesis that HARs function in human cells as regulatory sequences.

Recent pioneering work on the 3D structures of interphase chromosomes revealed specific, reproducible folding patterns of the chromosome fibers into spatially-segregated domain-like segments. In the mammalian nucleus, beads on a string linear strands of interphase chromatin fibers are folded into continuous megabase-sized topologically associating domains (TADs) that are readily detectable by the high-throughput analysis of interactions of chemically cross-linked chromatin (Dixon et al., 2012; Hou et al., 2012; Nora et al., 2012; Sexton et al., 2012). It has been hypothesized that TADs represent spatially-segregated neighborhoods of high local frequency of intrachromosomal contacts reflecting individual physical interactions between long-range enhancers and promoters of target genes in live cells (Dixon et al., 2012; Gorkin et al., 2014). Definition of TADs implies that neighboring TADs are separated by the sharp boundaries, across which the experimentally-detectable intrachromosomal contacts are relatively infrequent (Dixon et al., 2012; Gorkin et al., 2014). Genome-wide proximity placement analysis of 18,364 DNA sequences representing multiple diverse families of HSGRL within the context of the principal regulatory structures of the interphase chromatin in hESC identified TADs that are significantly enriched for HSGRL and termed rapidly-evolving in humans TADs (Glinsky, 2016). Concomitant HSGRL-associated increase of both the number and size of SEDs identifies the increasing of both regulatory complexity and functional precision of genomic regulatory networks due to convergence of TAD and SED architectures as one of the principal building blocks of the interphase chromatin structural changes during the evolution of *H. sapiens* (Glinsky, 2017). In this contribution, the analyses of human-specific features of pluripotency regulatory networks in hESC have been performed. The primary focus of this work was on the human-specific elements of the interphase chromatin architecture of specific SEDs and TADs responsible for transcriptional regulatory control in hESC of the *NANOG, POU5F1,* and *POU3F2* genes and comparative analyses of the adjacent four TADs of the extended *NANOG-*associated genomic region. Lastly, the putative mechanisms of the genome-wide regulatory effects of human-specific NANOG-binding sites (HSNBS) on expression of genes implicated in the fetal and adult brain development have been examined.

## Results

### Working models of *NANOG*, *POU5F1*, and *POU3F2* super-enhancers’ domains in hESC

One of the potential practical utilities of the analysis of HSGRL is the opportunity to use the available genomic information for building the experimentally testable working models of SEDs and TADs, which may be suitable for precise molecular and genetic definitions of critical structural elements of human-specific genomic regulatory networks. This notion was tested in the experiments designed to build working models of *NANOG*, *POU5F1*, and *POU3F2* super-enhancers’ domains in hESC (Fig. 1; Supplemental Fig. S1) and compare them to the corresponding SED structures recently identified in the mESC genomes (Dowen et al., 2014).

**Figure 1.**
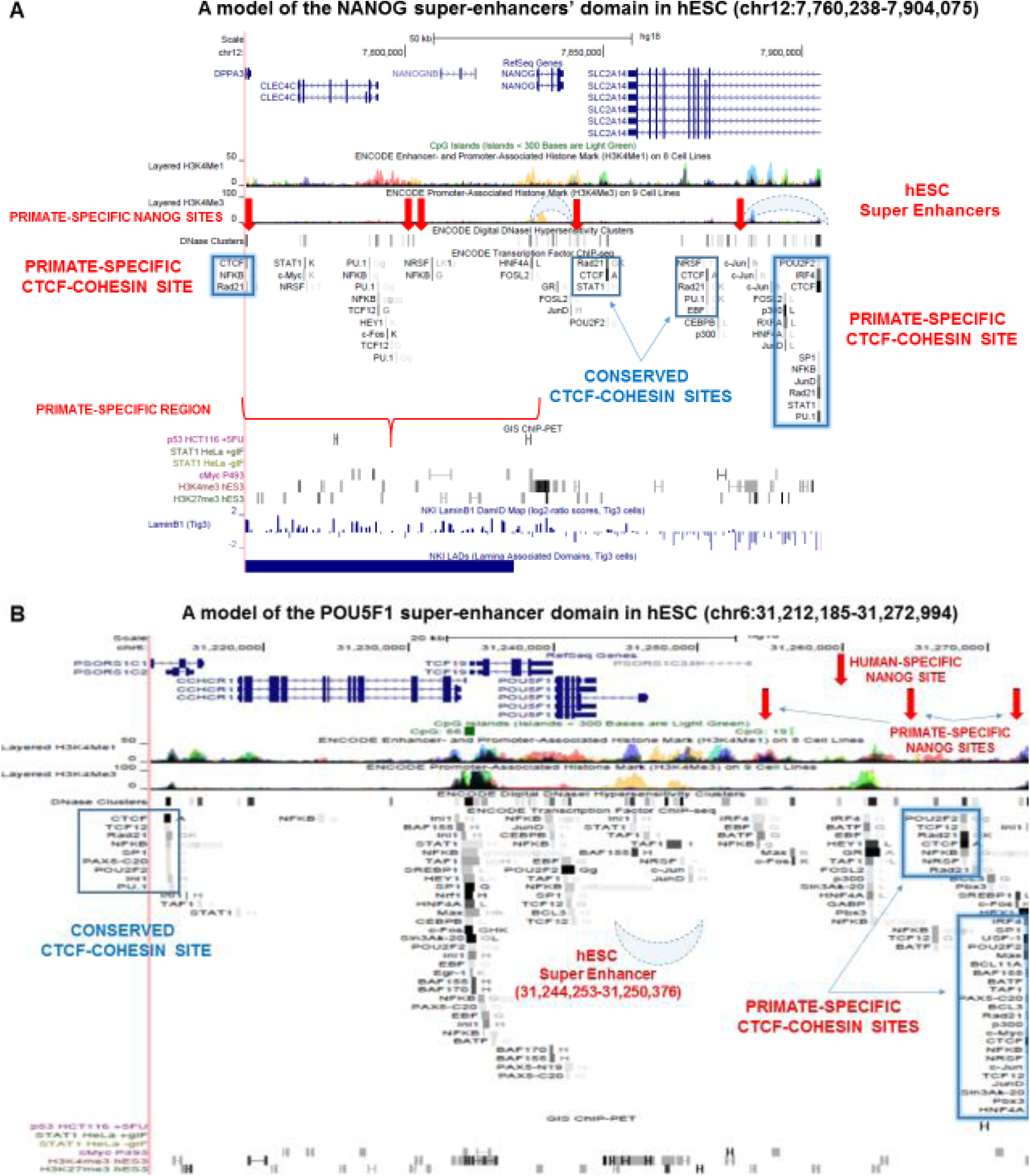
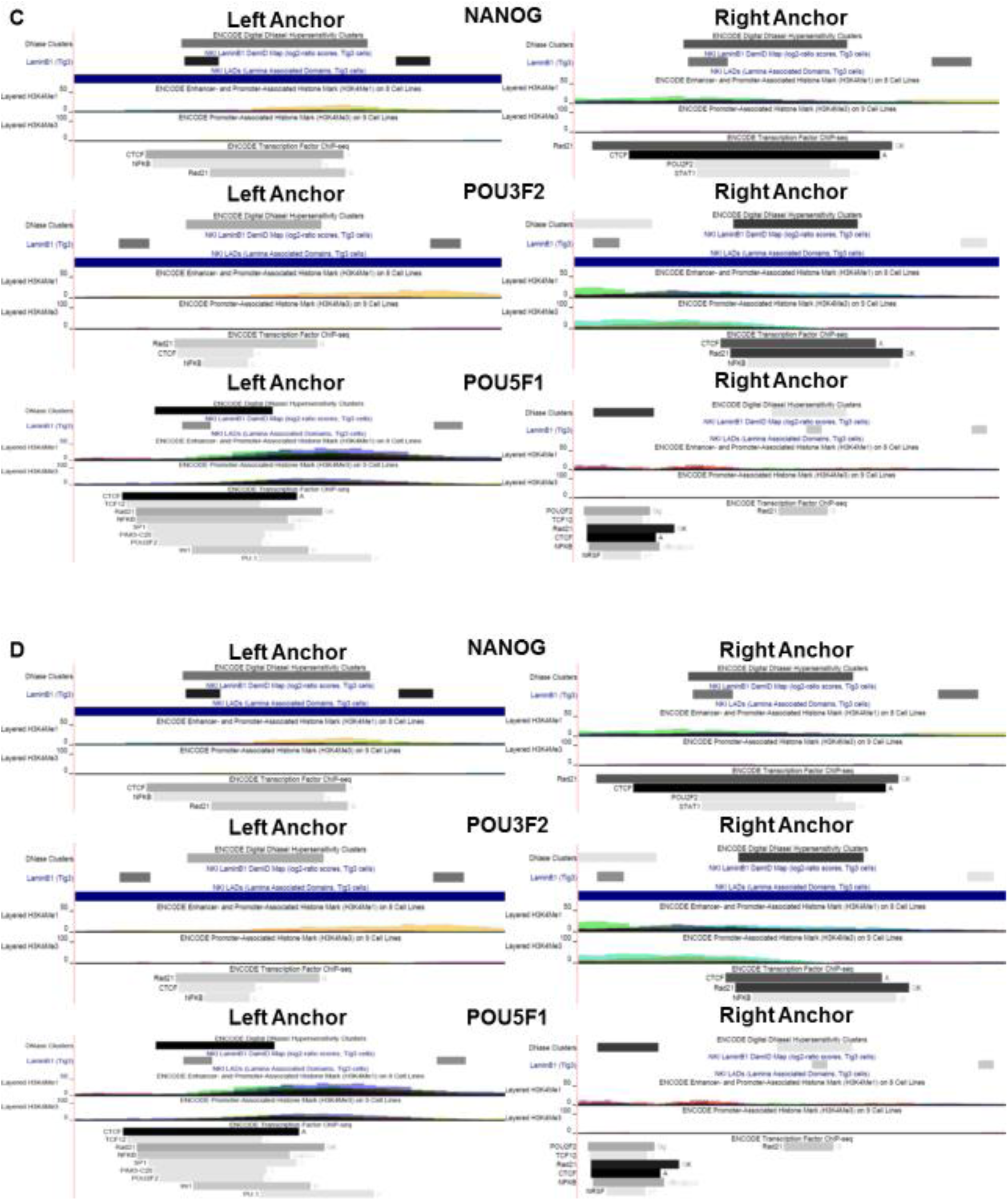
Genomic features of 2D models of individual super-enhancer domains (SEDs) in the hESC genome. A. UCSC Genome Browser view of the 2D model of the NANOG locus-associated super-enhancers’ domains in the hESC genome (chr12:7,760,238-7,904,075). Genomic coordinates of two hESC-super-enhancers are depicted by the semi-transparent arks; positions of 5 primate-specific NANOG-binding sites are marked by the red arrows; conserved and primate-specific overlapping CTCF/cohesin-binding sites that may be involved in formation of the anchors at the SED base are designated; genomic position of the primate-specific region harboring LAD sequence is depicted by the brace shape. Note that insertion of the primate-specific LAD sequence down-stream of the NANOG gene suggests that SED loop is more likely to be formed by folding of the upstream chromatin chain toward the nuclear lamina-bound DNA sequence (Fig. 2). B. A zoom-in view of the single super-enhancer model of the NANOG super-enhancer domain in hESC genome (chr12:7,760,238-7,851,119). Designations of symbols same as described in (A). C. UCSC Genome Browser view of the POU5F1 super-enhancer domain in the hESC genome (chr6:31,212,185-31,272,994). Genomic coordinates of the hESC-super-enhancer are depicted by the semi-transparent ark; positions of 3 primate-specific and one human-specific NANOG-binding sites are marked by the red arrows; conserved and primate-specific overlapping CTCF/cohesin-binding sites that may be involved in formation of the anchors at the SED base are designated. D. UCSC Genome Browser zoom-in views of the anchor regions at the bases of SEDs are shown. Note that SED anchor regions typically contain two LMNB1-binding sites near the overlapping CTCF/cohesin-binding sites, interactions of which secure the chromatin loop formation and segregation of insulated genomic neighborhoods of SEDs. These observations suggest that a spatial positioning of SED anchor regions is secured by the attachment to the nuclear lamina. A complete model of the POU3F2 super-enhancer domain in the hESC genome is reported in the Supplemental Fig. S1.

**Figure 2.**
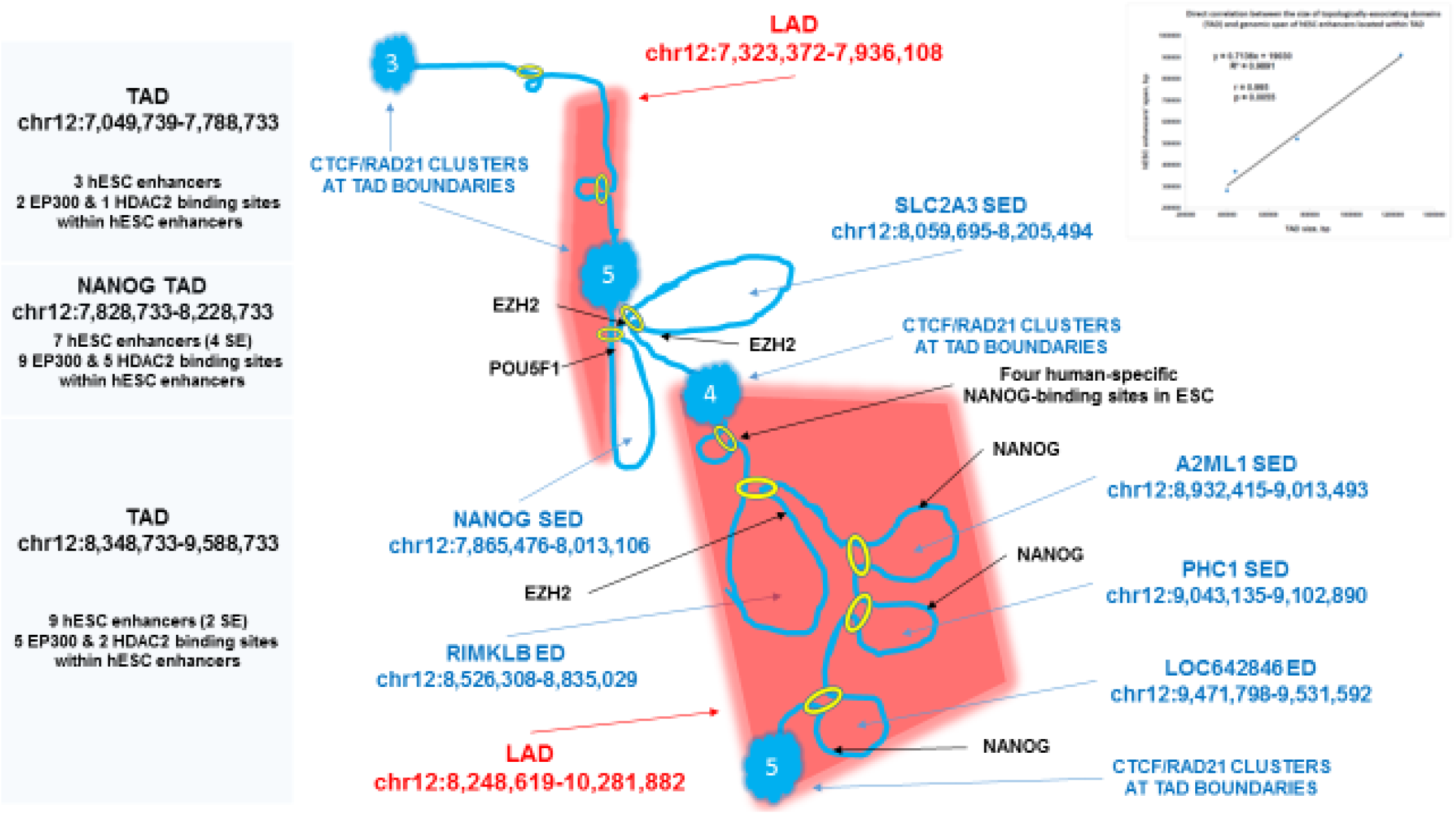
A snap-shot of principal genomic features of the 3D model of the interphase chromatin folding. A 3D model of spatial positioning of principal regulatory elements of the 2.54 Mb genomic region adjacent to the NANOG locus in the hESC genome is shown within the context of the 3 consecutive TADs and 4 boundary regions (depicted as blue clouds containing clusters of 3-5 overlapping CTCF/cohesin-binding sites). Positions of overlapping CTCF/cohesin-binding sites interactions of which secure the bases of chromatin loops are depicted by the yellow-colored donut shapes. Positions of key TFBS, including HSTFBS, are marked by the arrows. Attachment regions of blue-colored chromatin chains to the nuclear lamina are depicted by the placement of the corresponding segments within red-colored LADs. ED, hESC-enriched enhancers’ domain; SED, super-enhancers’ domains; EDs and SEDs are designated by the corresponding names of the protein-coding genes. Genomic coordinates of TADs, EDs, and SEDs are listed. Inset shows a highly significant direct correlation between the size of TADs and genomic span of hESC-enriched enhancers located within TADs (r = 0.995; p = 0.0055).

To identify the putative SED boundaries, a systematic search was conducted for nearest overlapping CTCF/cohesin sites located next to borders of the two hESC super-enhancers located within the TAD harboring the *NANOG* locus (Hnisz, et al., 2013). One of the notable features of the human *NANOG* SED is the presence of a large primate-specific region, which places the down-stream *NANOG* SED boundary within lamina-associated domain (LAD) harboring the primate-specific overlapping CTCF/cohesin site (Fig. 1A-B). Modeling of the human *NANOG* SED reveals that there are at least three possible designs of SED architecture that would incorporate both the *NANOG* SE and the target gene *NANOG* within the isolated genomic neighborhood of the looping structure formed as a result of interactions between two overlapping CTCF/cohesin sites (Fig. 1A). Two of SED design variants would include only one *NANOG* SE and would require the interactions between the primate-specific and conserved overlapping CTCF/cohesin sites enabling formation of the isolated genomic neighborhood loops of relatively smaller sizes (Fig. 1; Supplemental Fig. S1). The largest *NANOG* SED looping structure would result from interactions between two primate-specific overlapping CTCF/cohesin sites and it would incorporate both *NANOG* SEs within SED (Fig. 1). Notably, the postulated *NANOG* SED sequence harbors five primate-specific NANOG-binding sites previously identified in the hESC genome (Kunarso et al., 2010; Glinsky, 2015), which may indicate that NANOG protein exerts more efficient control over its own locus in primates’ genomes. In the mESC genome, the *NANOG* SED is markedly smaller occupying just 52 Kb and harboring only a single *NANOG* SE (Dowen et al., 2013). The entire SED region down-stream of the *NANOG* gene, which contain the *NANOG* SE sequence, has no orthologous sequences in primate and human genomes and appears replaced by the large primate-specific sequence in the human genome (Fig. 1; Supplemental Fig. S1).

Similarly dramatic rearrangements of the SED architecture in the hESC genome are apparent when additional examples of the SED structures insulating SEs regulating *POU5F1* and *POU3F2* target genes were analyzed (Fig. 1C; Supplemental Fig. S1). In all instances, the genomic looping structure-enabling interactions between two overlapping CTCF/cohesin sites would require the involvement of at least one primate-specific overlapping CTCF/cohesin-binding site. The *POU5F1* SED region harbors one human-specific and three primate-specific NANOG-binding sites (Fig. 1C; Supplemental Fig. S1), whereas the *POU3F2* SED contains ten hESC-enhancers and twenty-eight primate-specific TFBS for NANOG (15 sites), POU5F1 (6 sites), and CTCF (7 sites) proteins (Supplemental Fig. S1). The example of *POU3F2* SED is of particular interest because it is located within one of the rapidly-evolving in humans TADs and, according to recent experiments, the *POU3F2* gene plays a pivotal role in the highly efficient conversion of human pluripotent stem cells as well as non-neural human somatic cells into functional neurons (Pang et al., 2011; Wapinski et al., 2013). Of note, sequences near the overlapping CTCF/cohesin sites, interactions between which bring about the SED structures, appear to have a common structural element comprising two conserved in individual human genomes LMNB1-binding sites adjacent to CTCF/cohesin sites (Fig. 1D; Supplemental Fig. S1). These LMNB1-binding sites may serve as anchors to nuclear lamina for DNA sequences harboring the overlapping CTCF/cohesin sites, thus thermodynamically enhancing the likelihood of creation and/or stabilizing the maintenance of the base of SED structures.

Applying similar strategy as a guidance to infer the putative SED and TAD structures, it is possible to build the working models of the multidimensional chromatin architecture for large genomic regions spanning 2.54-3.26 Mb and containing 3-4 TAD structures (Figs. 2 & 3; Supplemental Fig. S2). Key elements of this design strategy entails the following considerations:

i. Defining the positions of the SED base by the overlapping CTCF/cohesin sites interacting with each other to create the SED looping structure;
ii. Mapping the putative bending sites of linear chromatin fibers based on the positions of Alu clusters near the overlapping CTCF/cohesin sites;
iii. Requiring that SEs and their target genes are located within the same insulated chromatin looping structure;
iv. Taking into account the relative strength of the chromatin fiber interactions with nuclear lamina based on placements of corresponding DNA sequences within or outside LADs and/or based on the quantity of LMNB1-binding sites within a region;
v. Strictly adhering to the previously defined genomic positions of TAD and LAD boundaries, ESC enhancers, primate-specific TFBS, and HSGRLs.

**Figure 3.**
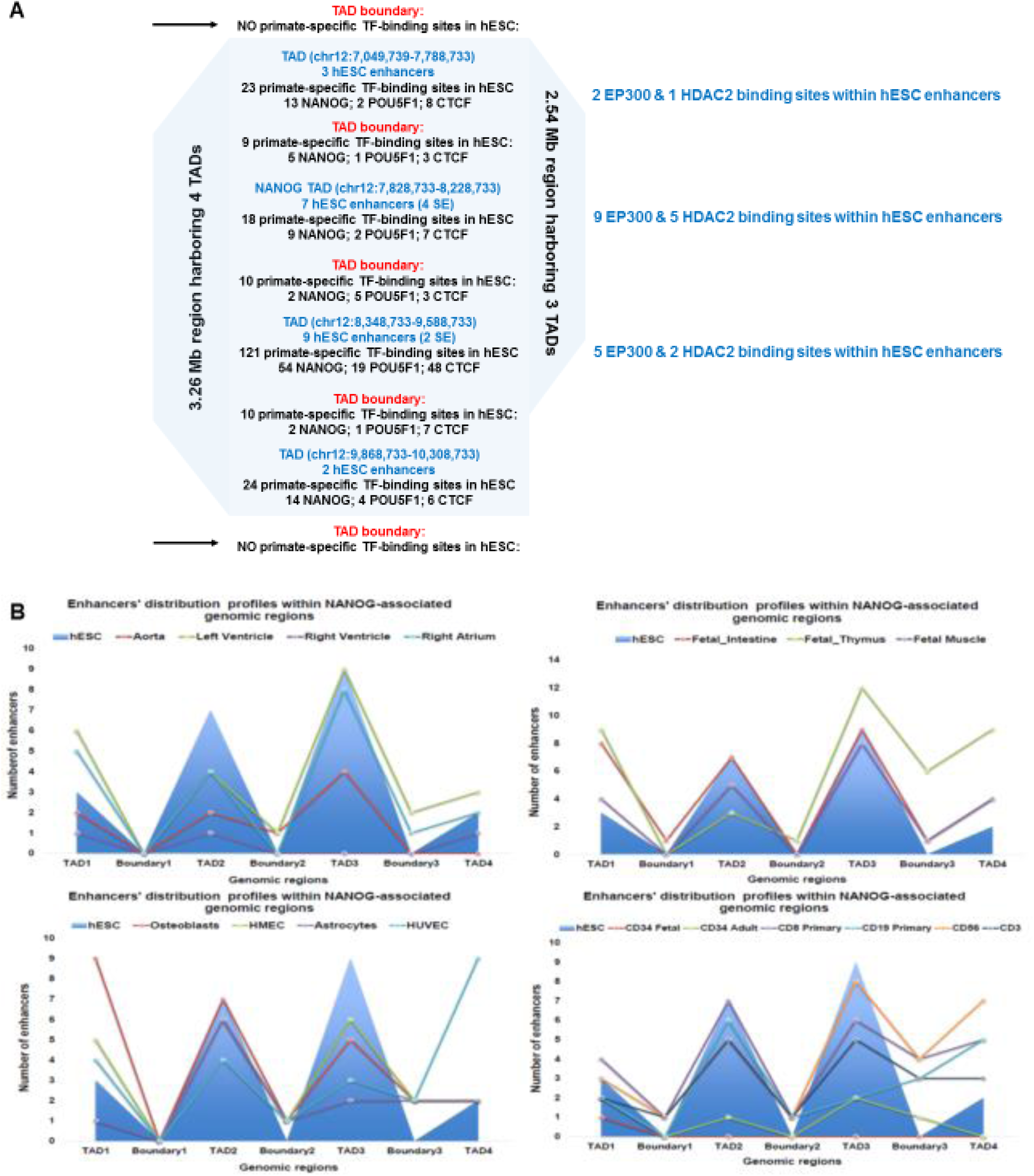
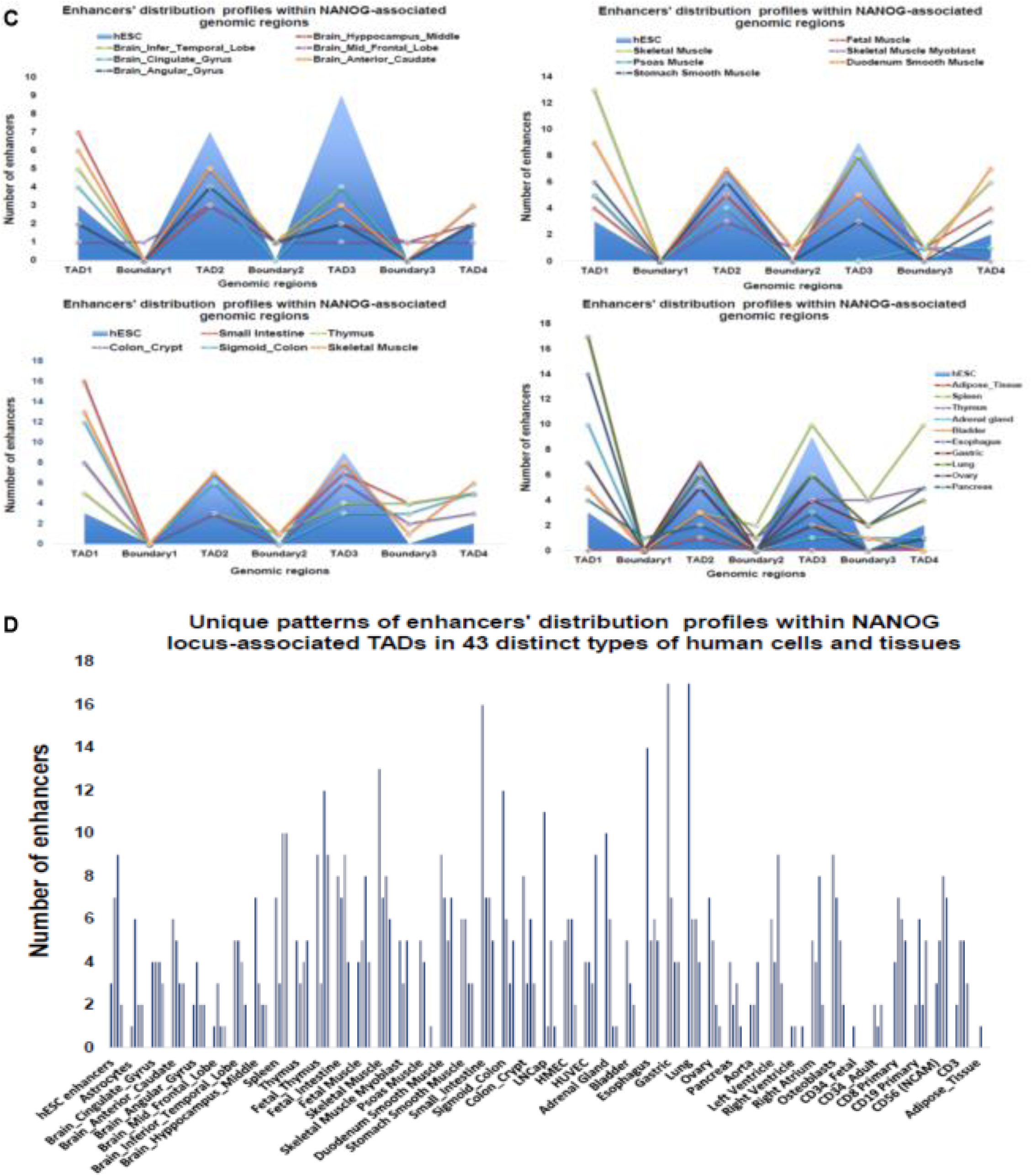
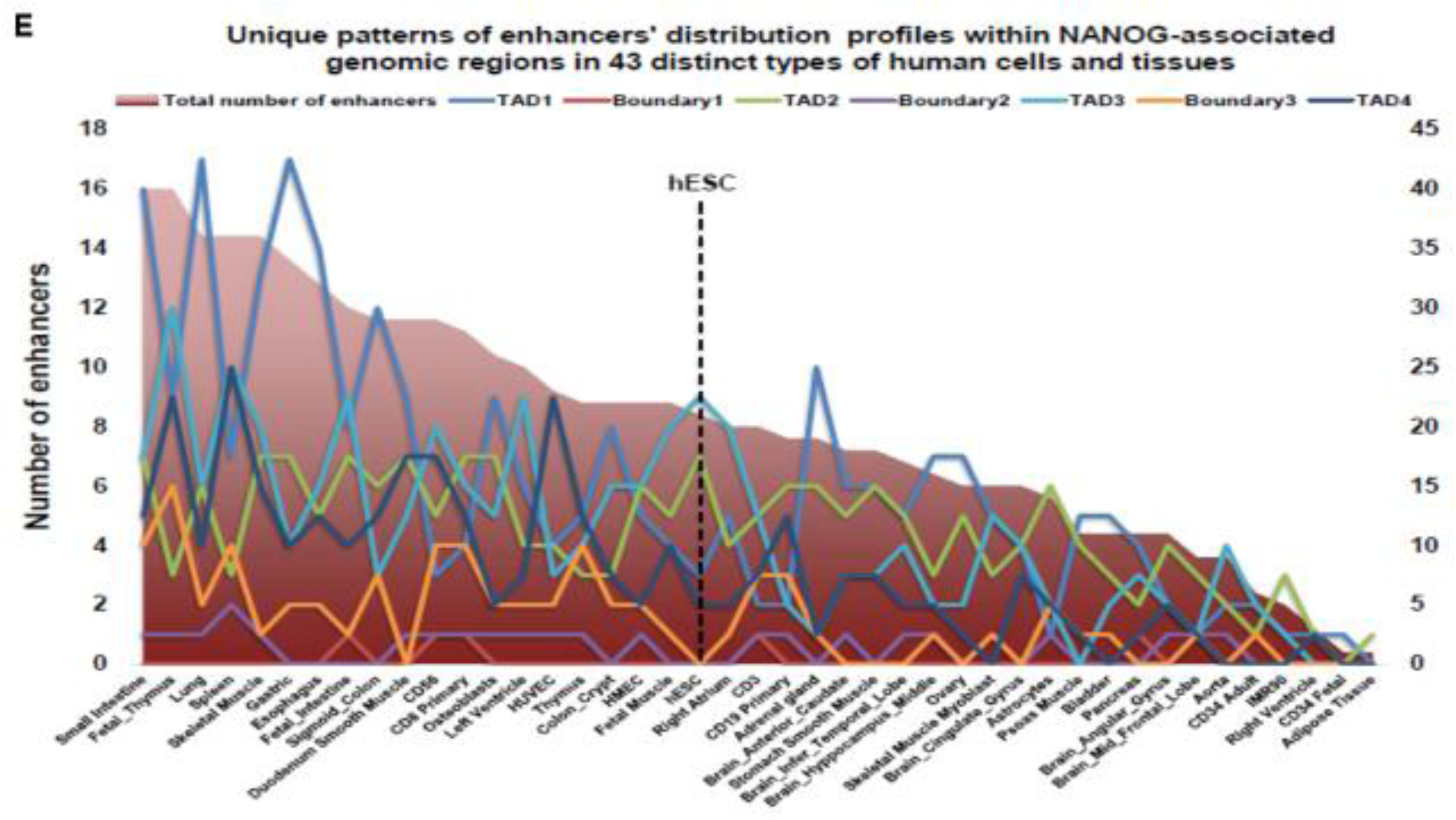
Conserved profiles and uniquely distinct patterns of H3K27ac peaks’ distribution within the NANOG locus-associated genomic region in 43 types of human cells and tissues. Regional genomic maps of H3K27ac peaks/SE distributions depicting numbers of SEs placed within 3.26 Mb genomic region containing four TADs and three TAD boundaries (A) for individual types of human cells and tissues against the background of SE distribution profiles within the same region in the hESC genome. Note that all individual profiles of SE distributions in human cells and tissues (B, C), which reflect distribution profiles of H3K27ac peaks, follow the general pattern of SE distributions established in the hESC genome (figures B & C). One of the notable common elements that emerged in many differentiated cells and tissues is the appearance of SEs within the genomic regions demarcating TAD boundaries in the hESC genome, suggesting that TAD boundaries and TAD structures undergo visible rearrangements in differentiated cells and tissues. Erasing the hESC TAD boundaries during differentiation due to appearance of new cell-type specific SEs is consistent with observed decreasing numbers of TADs in differentiated cells and tissues (figure 4). D, E. Unique patterns of H3K27peaks’/enhancers’ distribution profiles in 43 types of human cells and tissues. In figure E, the data set was sorted in descending order of total numbers of enhancers (right Y axis) located within the region in each cell and tissue type. The position of hESC within this distribution is shown by the dotted line. The names of cells and tissues are listed. See text for details.

**Figure 4.**
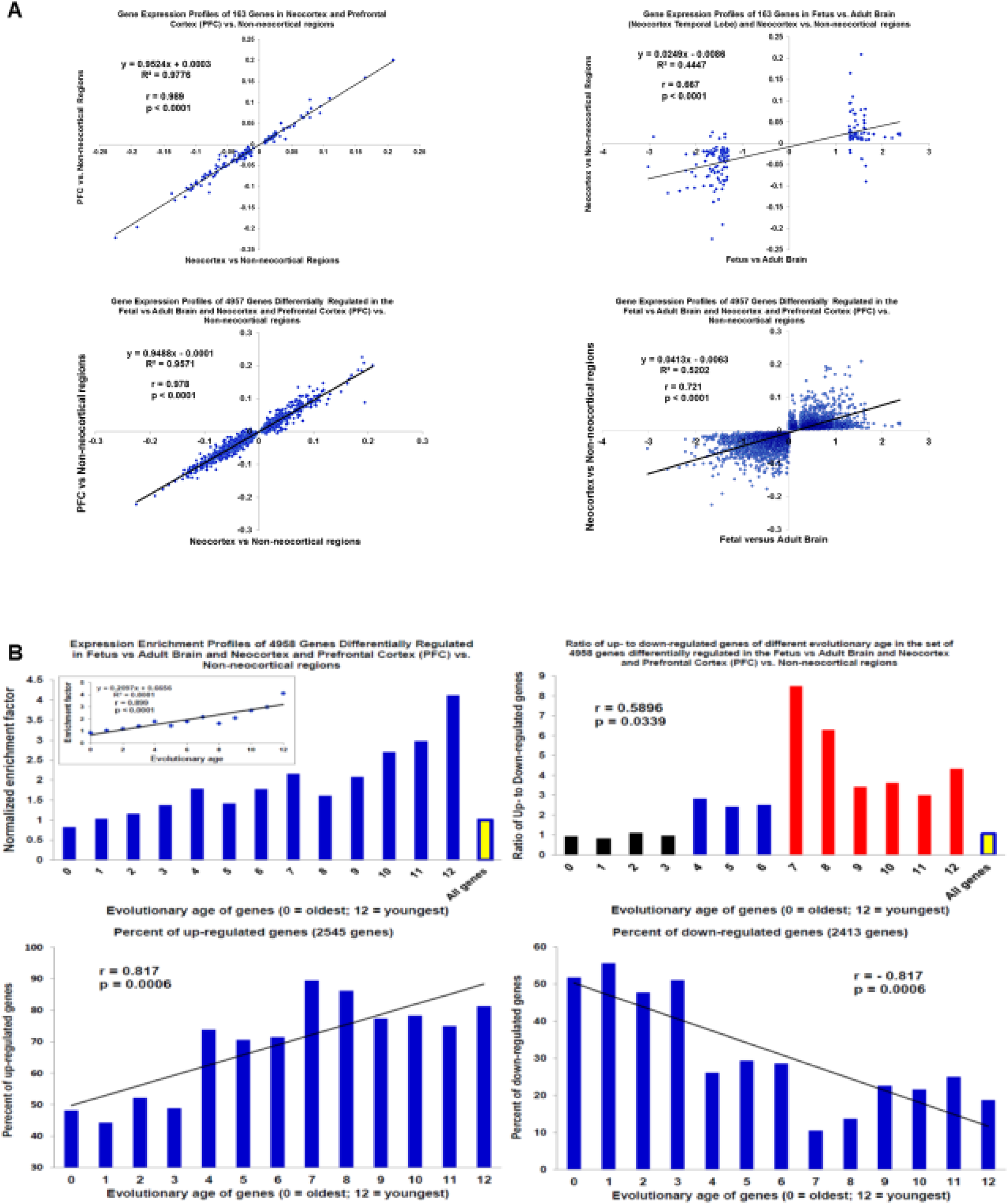
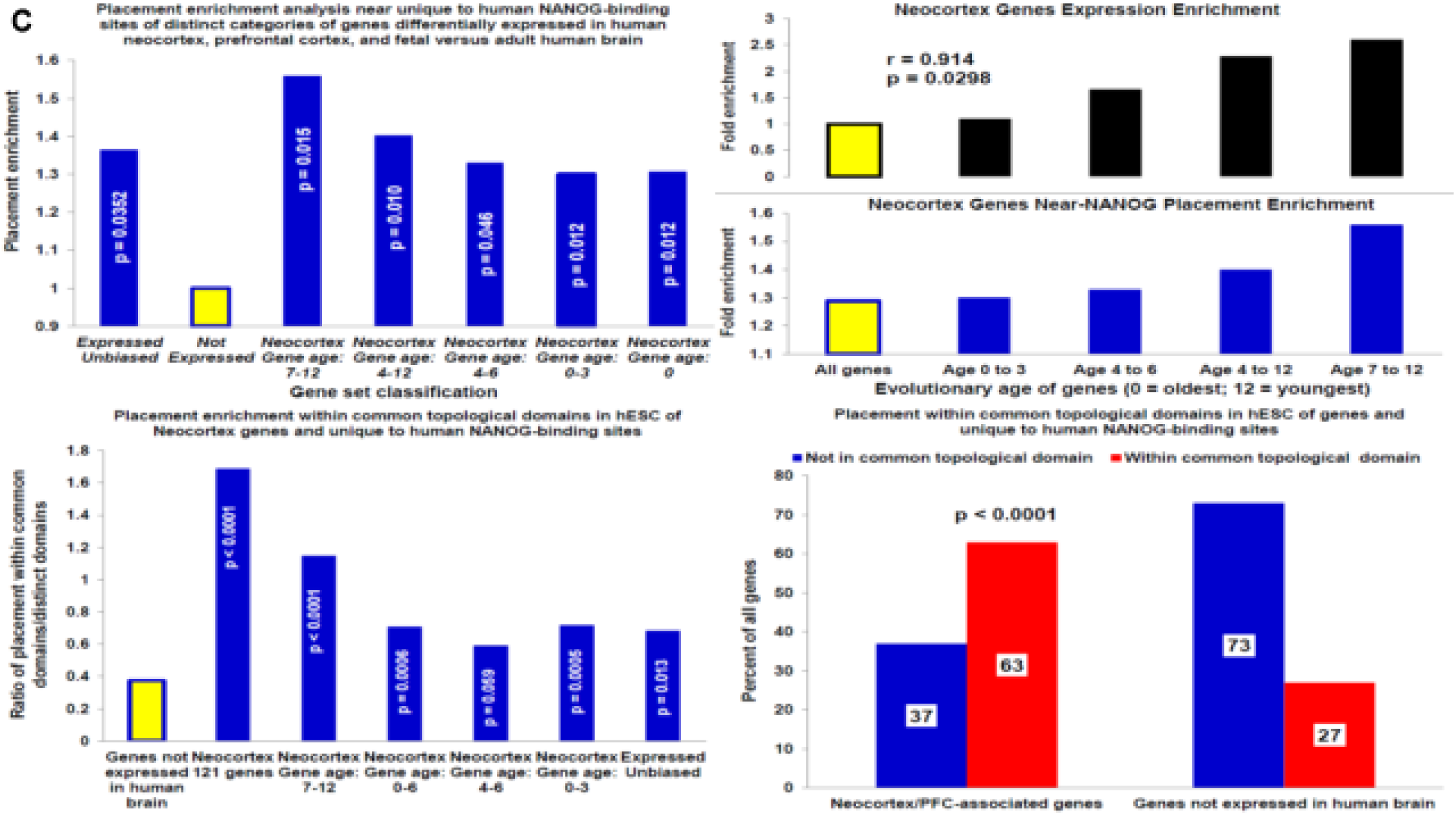
Identification and characterization of the 4,957 genes’ expression signature in the human fetal neocortex temporal lobe retaining the expression changes acquired in the embryo for many years of human brain development and maintaining highly concordant expression profiles in the neocortex and prefrontal cortex regions of adult human brain. A. Concordant expression profiles of 163 genes with at least 20-fold expression changes in the fetal neocortex temporal lobe that also manifest significant expression changes in the neocortex versus non-neocortical regions of adult human brain (top right panel). Note that expression changes of these 163 genes are highly concordant in both neocortex and prefrontal cortex regions of adult human brain versus non-neocortical regions (top left panel). Bottom two panels show highly concordant expression profiles of 4,957 genes with significant expression changes in fetal versus adult human brain and neocortex versus non-neocortical regions of adult human brain (bottom right panel) as well as either prefrontal cortex or neocortex versus non-neocortical regions of adult human brain (bottom left panel). B. Analysis of expression enrichment patterns of 4,957 genes suggests a bias toward up-regulation of evolutionary young genes and down-regulation of evolutionary old genes. See text for detail. C. Placement enrichment analysis of neocortex/prefrontal cortex signature genes near human-specific NANOG-binding sites (HSNBS) suggests a bias toward evolutionary young coding genes and genes expressed in brain compared with genes that are not expressed in human brain. Bottom right panel shows enrichment of co-localization within common TADs for neocortex/prefrontal cortex signature genes and HSNBS compared with genes that are not expressed in human brain.

Using this strategy, a model has been built representing a global view of multidimensional chromatin folding patterns within the 2.54 Mb genomic region containing the *NANOG*-residing TAD and two adjacent neighboring TADs (Fig. 2). A more detailed depiction of sequential steps of the model building is presented in the Supplemental Fig. S2. It seems evident, that one of the principal structural elements defining directionality of the *NANOG* SED looping structure in the human genome is the insertion immediately downstream of the *NANOG* gene of the primate-specific sequence, which is tightly attached to the nuclear lamina as depicted in the Fig. 2 by placement of the corresponding part of the *NANOG* SED within red-colored LAD. The remaining part of the *NANOG*-harboring TAD represents the genomic region located upstream of the *NANOG* SED. It is not attached to the nuclear lamina and contains the *SLC2A3* SED, which is located in its entirety outside of the LAD boundaries (Fig. 2). Further upstream of the *SLC2A3* SED at the boundary region between the *NANOG*-harboring TAD and adjacent upstream TAD, the chromatin fiber re-attaches to nuclear lamina and places within LAD the entire TAD containing 4-5 insulated neighborhood looping structures (Fig. 2). It will be of interest to determine how transitions of placements of chromatin fibers between positions within and outside LAD would affect the creation of SED structures and performance of insulated within SED genomic regulatory circuitry. One of the near-term practical applications of this approach to multidimensional modeling of SED and TAD structures became apparent when the regulatory elements residing within corresponding TADs and sub-TAD structures were catalogued and reviewed.

### Conserved profiles and uniquely distinct patterns of H3K27ac peaks’ distribution within the *NANOG*-associated genomic region in 43 types of human cells and tissues

One of the characteristic regulatory features systematically associated with hESC enhancers located within four TADs of the *NANOG* region is the presence of multiple binding sites for both EP300 histone acetyltransferase and histone deacetylase 2, HDAC2 (Figs. 2 & 3; Supplemental Fig. S2). Both histone acetyltransferases (HATs) and histone deacetylases (HDACs) have essential and pleiotropic roles in regulation of stem cell self-renewal by maintaining expression of master TFs regulating the pluripotent state and controlling the core transcriptional regulatory networks of embryonic stem cells and their differentiated progenies (Chen et al., 2008; Fazzio et al., 2008; Zhong X, Jin Y, 2009; Li et al., 2012; Jamaladdin et al., 2014). These observations are likely reflect the requirement for constitutive activities of both HATs and HDACs at transcriptionally active regulatory elements, because nucleosome maintenance and turnover are highly dynamic at these loci due to markedly accelerated exchange rates of histones, which appear directly associated with the histone modifications defining the active chromatin regulatory state. For example, H3.3-containing nucleosomes undergo rapid turnover at active enhancers and promoters (Kraushaar et al., 2013). This fast turnover is positively correlated with active chromatin histone modification marks, including H3K4me1, H3K4me3, H3K9ac, and H3K27ac (Kraushaar et al., 2013). In contrast, the nucleosome exchange rate is negatively correlated with repressive chromatin marks H3K27me3 and H3K9me2 at regulatory loci and H3K36me3 modifications within gene bodies (Kraushaar et al., 2013).

It is intriguing that binding sites for HDAC2 but not HDAC1 were detected in this region. Of note, it has been demonstrated that a single allele of *Hdac2* but not *Hdac1* is sufficient to rescue normal mouse brain development in double knockout *Hdac1*^-/-^*Hdac2*^-/-^ mice and that HDAC2 has a unique indispensable role in controlling the fate of neural progenitor cells (Hagelkruys et al., 2014). Forced neuron-specific overexpression of HDAC2, but not HDAC1, reduced dendritic spine density, synapse number, synaptic plasticity, and memory formation in mice (Guan et al., 2009). Consistently, reduction of synapse number and learning impairment of *Hdac2*-overexpressing mice were ameliorated by chronic HDACi treatment (Guan et al., 2009). In the mouse models of neurodegeneration and in brains of patients with Alzheimer’s disease, cognitive capacities were severely impaired by the epigenetic blockade of gene transcription at specific genetic loci important for learning and memory, which is mediated by the increased HDAC2 activity at these sites (Gräff et al., 2012). Neurodegeneration-associated memory impairments, abnormal structural and synaptic plasticity, and diminished expression of genes regulating learning and memory were reversed following shRNA-mediated knockdown of HDAC2 overexpression (Gräff et al., 2012). Critical role of HDAC2 in regulation of brain functions is supported by the recent experiments on individual neurons demonstrating that HDAC2 cell autonomously suppresses excitatory activity and enhances inhibitory synaptic function in CA1 pyramidal neurons (Hanson et al., 2013).

Since SE structures in genomes of human cells were defined based on the genomic profiles of H3K27ac peaks (Hnisz et al., 2013), it is reasonable to argue that patterns of SE placements within TAD reflect genomic positions of H3K27ac peaks and should faithfully represent the snap-shot of the balance of continuing HAT and HDAC activities at specific regulatory sites within a region. Based on these considerations, genomic maps of SE distributions were drawn for 43 distinct types of human cells and tissues by retrieving genomic coordinates of SEs from the recently reported comprehensive catalogue of SEs in human body (Hnisz et al., 2013). Regional genomic maps of SE distributions shown in Figs. 2 & 3 depict numbers of SEs placed within 3.26 Mb genomic region containing four TADs and three TAD boundaries for individual types of human cells and tissues against the background of SE distribution profiles within the same region in the hESC genome. It is evident that all individual profiles of SE distributions in human cells and tissues, which reflect distribution profiles of H3K27ac peaks, follow the general pattern of SE distributions established in the hESC genome (Figs. 3B & 3C). One of the notable common elements that emerged in many differentiated cells and tissues is the appearance of SEs within the genomic regions demarcating TAD boundaries in the hESC genome, suggesting that TAD boundaries and TAD structures underwent visible rearrangements in humans’ differentiated cells and tissues compared with hESC. Erasing the hESC TAD boundaries during differentiation due to appearance of new cell-type specific SEs is consistent with observed decreasing numbers of TADs in differentiated cells and tissues (Glinsky, 2017).

Transformation of these data into different types of charts for patterns’ visualization (Figs. 3B-3E) revealed that all analyzed human cells and tissues manifest unique profiles of SE/H3K27ac distributions within the 3.26 Mb *NANOG*-associated genomic region. It seems quite remarkable that quantitative distribution profiles of the very small set of markers located within ~0.1% of human genome appear to distinguish 43 distinct types of human cells and tissues. This notion is best illustrated in the Fig. 3E showing the data set of 43 distinct types of human cells and tissues that were sorted in descending order based on total numbers of SEs located within the 3.26 Mb *NANOG*-associated genomic region. It is clearly visible that most samples are easily distinguishable by the total number of SEs located within the region and samples that have the same numbers of SE’s manifest readily distinct SE/H3K27ac distribution profiles (Fig. 3E). These results seem to point to a stochastic, i.e., random and probabilistic, nature of the underlying causal mechanisms driving the emergence of unique profiles of SE/H3K27ac distributions in differentiated cells and tissues from the singular parental SE/H3K27ac distribution profile in hESC. Consistent with this hypothesis, 45.2% of human cells’ and tissues’ samples have the higher numbers of SE’s placements compared with hESC, whereas the SE’s placement numbers are lower compared with hESC in 54.7% of samples (Fig. 3E). Collectively, this analysis appears to indicate that the emergence of unique cell type-specific patterns of SE/H3K27ac placements may have been triggered by the small bidirectional changes of the balance between the HAT’s and HDAC’s activities in the genome of hESC that evolved during the differentiation process into arrays of quantitatively distinct patterns of SE/H3K27ac distributions in different types of cells and tissues in the human body.

### Putative regulatory consequences of targeted placements of HSGRL affecting expression of protein-coding genes

Observed increasing regulatory complexity in human genomes associated with targeted placements of HSGRL is consistent with a view that increasing genomic complexity within fixed environments represents a major evolutionary trend in the natural world (Adami et al., 2000). Further analysis suggests that these changes of nuclear regulatory architecture may facilitate the enhanced precision of regulatory interactions between enhancers and target genes within insulated neighborhoods of chromatin regulatory networks. The boundaries of a majority of TADs are shared by the different cell types within an organism, segregating genomes into distinct regulatory units that harbor approximately seven protein-coding genes per TAD (Dixon et al., 2012; Smallwood and Ren, 2013). In contrast, nearly two-third of revTADs (39 of 60 revTADs; 65%) contain five or less protein-coding genes. Among these, eight revTADs harbor just one protein-coding gene and two revTADs contain only non-coding RNA genes. Of the twenty one revTADs harboring more than five protein-coding genes, seven revTADs contain clusters of functionally-related protein-coding genes: 3 revTADS on chr11 harbor 61 genes encoding olfactory receptors; 3 revTADs on chr19 harbor 31 genes encoding zinc finger proteins; and one revTAD on chr20 contains eight beta-defensins’ genes. Importantly, the expression of genes residing within the same TAD appears somewhat correlated and some TADs tend to contain predominantly actively transcribed genes while others have mostly repressed genes (Cavalli and Misteli, 2013; Gibcus and Dekker, 2013; Nora et al., 2012). Collectively, these results suggest that observed enrichment of HSGRL placements in specific genomic regions may facilitate the enhanced precision of regulatory functions in human genomes by targeting TADs harboring relatively few protein-coding genes and/or TADs containing clusters of functionally-related protein-coding genes, thus preferentially affecting insulated regulatory networks governing selected key developmental events.

Recently reported results of proximity placement analyses of human-specific NANOG-binding sites (HSNBS) revealed their associations with coding genes governing physiological development and functions of nervous and cardiovascular systems, embryonic development, behavior, as well as development of a diverse spectrum of pathological conditions (Glinsky, 2015). Here this approach was applied to dissect the expression profiles of protein-coding genes implicated in the development of the fetal and adult brain of *H. sapiens* (Johnson et al., 2009; Zhang et al., 2011) and examine their co-localization patterns with HSNBS in the human genome. During the first step of this analysis, a set of 251 genes that are at least 20-fold up- or down-regulated in the neocortex temporal lobe of the fetal versus adult human brain has been identified. This set of 251 genes was designated a gene expression signature of the human fetal neocortex temporal lobe. It was of interest to ascertain the expression changes of these genes, if any, in the neocortex, the newest part of human brain, and in the prefrontal complex, which is implicated in complex cognitive behavior, compared with non-neocortical regions in both fetal and adult human brains. Expression of two-third of genes comprising the 251-gene expression signature of the human fetal neocortex temporal lobe is significantly different in the neocortex compared with non-neocortical regions of adult human brain and gene expression profiles of these genes in neocortex and prefrontal cortex are highly concordant (Fig. 4). Of note, 78% of genes differentially regulated in the neocortex versus non-neocortical regions manifest significant changes of expression in the prefrontal cortex versus non-neocortical regions of human brain as well. Remarkably, 92.6% of genes differentially regulated in both fetal versus adult brain and neocortex versus non-neocortical regions exhibit the same direction of expression changes, that is genes that are up-regulated in the fetal brain remain up-regulated in the neocortex and genes that are down-regulated in the fetal brain remain down-regulated in the neocortex (Fig. 4). Thus, a majority of genes (60%) that acquired most dramatic expression changes in the human fetal neocortex temporal lobe appears to retain for many years the significance of expression changes acquired in the embryo and maintain highly concordant expression profiles in the neocortex and prefrontal cortex regions of adult human brain. Investigation of the placement enrichment pattern of HSNBS located near these neocortex/prefrontal cortex-associated genes revealed the most significant enrichment of HSNBS placement at the genomic distances less than 1.5 Mb with a sharp peak of the enrichment p value at the distance of 1.5 Mb (Supplemental Fig. S3), suggesting that HSNBS may play a role in regulation of expression of these genes during embryogenesis.

These observations prompted the detailed analysis of all 12,885 genes, expression of which is significantly different in the human fetal neocortex temporal lobe compared with the adult brain (Johnson et al., 2009; Zhang et al., 2011), and examine their expression in the neocortex and prefrontal cortex regions of the adult human brain. Remarkably, the analysis of these fetal neocortex temporal lobe-associate genes revealed that expression of 4,957 genes is significantly different in the neocortex compared with non-neocortical regions of adult human brain and gene expression profiles of these genes in neocortex and prefrontal cortex are highly concordant (Fig. 4). These results demonstrate that a very large number of genes, consisting of ~20-25% of all protein-coding genes in the human genome, acquires in embryo significant expression changes, which are stably maintained throughout the embryogenesis, infancy, and the adulthood as a highly concordant gene expression profile in the human fetal neocortex temporal lobe as well as in the neocortex and prefrontal cortex regions of adult human brain. It provides a compelling argument that gene expression profiles characteristic of the critical brain regions, which are essential for development of unique to human cognitive and behavioral functions, are established to a significant degree during the embryogenesis, retained for many years of human development, and maintained in the adulthood. In agreement with this idea, there is an increased enrichment of 5hmC at intragenic regions that are already hyper-hydroxymethylated at the fetal stage during human frontal cortex development, demonstrating that adult patterns of genic 5hmC in the frontal cortex are already evident in the immature fetal brain (Lister et al., 2013). In the human brain, transcriptional activity is associated with intragenic 5hmC enrichment and adult 5hmC patterns for cell type-specific genes appear established *in utero*; in contrast, loss of 5hmC enrichment is associated with developmentally coupled transcriptional down regulation of gene expression (Lister et al., 2013).

Recent experiments discovered that that there is a strikingly accelerated recruitment of new, evolutionary young genes during the early development of human brain leading to the increased expression of a significantly larger proportion of young genes in fetal or infant brains of humans compared with mouse (Zhang et al., 2011). Next, the detailed analysis of the evolutionary age of genes comprising the 4,957-gene expression signature of the neocortex/prefrontal cortex regions of human brain was carried-out. To this end, all 4,957 genes were segregated into thirteen sub-groups based on their respective evolutionary age, ranging from 0 (oldest genes) to 12 (youngest genes) as previously defined by Zhang et al. (2011). The gene expression enrichment factors were calculated for each individual evolutionary age sub-group by comparisons of corresponding gene age-associated distribution metrics, which were derived from the analysis of gene age-associated distribution profiles of all 19,335 genes interrogated in gene expression profiling experiments and a set of 12,885 genes with expression changes significantly different in fetal versus adult brain. The resulting values for each evolutionary age sub-group of genes were normalized to the corresponding numerical values obtained for all genes within the corresponding set. The results of these analyses revealed that the relative enrichment factors of evolutionary young genes are higher compared to evolutionary older counterparts, which appears to be associated exclusively with the fraction of up-regulated genes of the 4,957-gene signature of the neocortex/prefrontal cortex regions of human brain (Fig. 4).

Next, the placement enrichment analysis of HSNBS located in close proximity to the genomic coordinates of the 4,957 genes was performed. In these experiments, the estimates of two different placement enrichment metrics were computed. The numerical values of the first metric is based on the quantitation of genes residing within the genomic distance of 1.5 Mb or less from the nearest HSNBS (Fig. 4). The threshold of 1.5 Mb was chosen based on the quantitative definition of the genomic distance associated with the most statistically significant enrichment of HSNBS placement near neocortex/prefrontal cortex associated genes (Supplemental Fig. S3). The numerical values of the second metric is based on the quantitation of genes residing within and outside of the common TAD with HSNBS (Fig. 4). In both experimental settings, the corresponding placement enrichment values for each sub-group of genes were compared to the values calculated for two control gene sets: 447 genes that are not expressed in the human brain and 357 expressed unbiased genes, i.e., genes that are uniformly expressed in human brain and do not manifest statistically significant differences of expression. Results of these analyses were highly consistent regardless of the utilized placement enrichment metrics: it appears that placements of HSNBS favors the evolutionary younger genes of sub-groups 7-12 compared with the evolutionary older genes or genes that are not expressed in the human brain (Fig. 4). It is important to mention that all genes comprising both control gene sets were assigned to the evolutionary age groups 7-12, thus excluding the possibility that placement enrichment differences are due to genes of the control gene sets being belong to the evolutionary older genes.

Collectively, these data are highly consistent with the idea that HSNBS may have contributed to the establishment during the embryogenesis of the genome-wide expression changes characteristic of neocortex and prefrontal cortex regions of human brain, which are retained as a highly concordant 4,957-gene expression signature for many years of human development and maintained in the adulthood. The validity of this idea was further explored utilizing the Ingenuity Pathway Analysis software (http://www.ingenuity.com/) to identify and analyze possible developmental and pathophysiological regulatory networks comprising of HSNBS-associated genes, which appear implicated in development of human neocortex and prefrontal cortex regions. The Ingenuity software identified two main candidate regulatory networks and predicted five potential top regulators and eight their immediate target genes (Fig. 5). Of note, seven of eight immediate target genes of the top five putative upstream regulators were also identified as regulatory elements of the two main networks, which are marked by the five-pointed stars in the Fig. 5. One exception is the *INSM1* and *NEUROD1* genes that appear to form an interconnected and highly biologically significant axis, which plays a crucial role in maintaining mature pancreatic β-cell functions via cooperating interactions of *INSM1*, *NEUROD1*, and *FOXA2* genes and combinatorial binding of INSM1, NEUROD1, and FOXA2 proteins to regulatory DNA sequences (Jia et al., 2015). Furthermore, the *Insm1* gene is required for proper differentiation of all types of endocrine cells in the anterior pituitary gland, including pituitary cells producing thyroid-stimulating hormone, follicle-stimulating hormone, melanocyte-stimulating hormone, adrenocorticotrope hormone, growth hormone and prolactin (Welcker et al., 2013). Because it has been previously demonstrated that *Insm1* gene is required for development and differentiation of endocrine cells in the pancreas, intestine and adrenal gland (Gierl et al., 2006; Wildner et al., 2008), it has been defined as the essential pan-endocrine transcription factor (Welcker et al. (2013).

**Figure 5.**
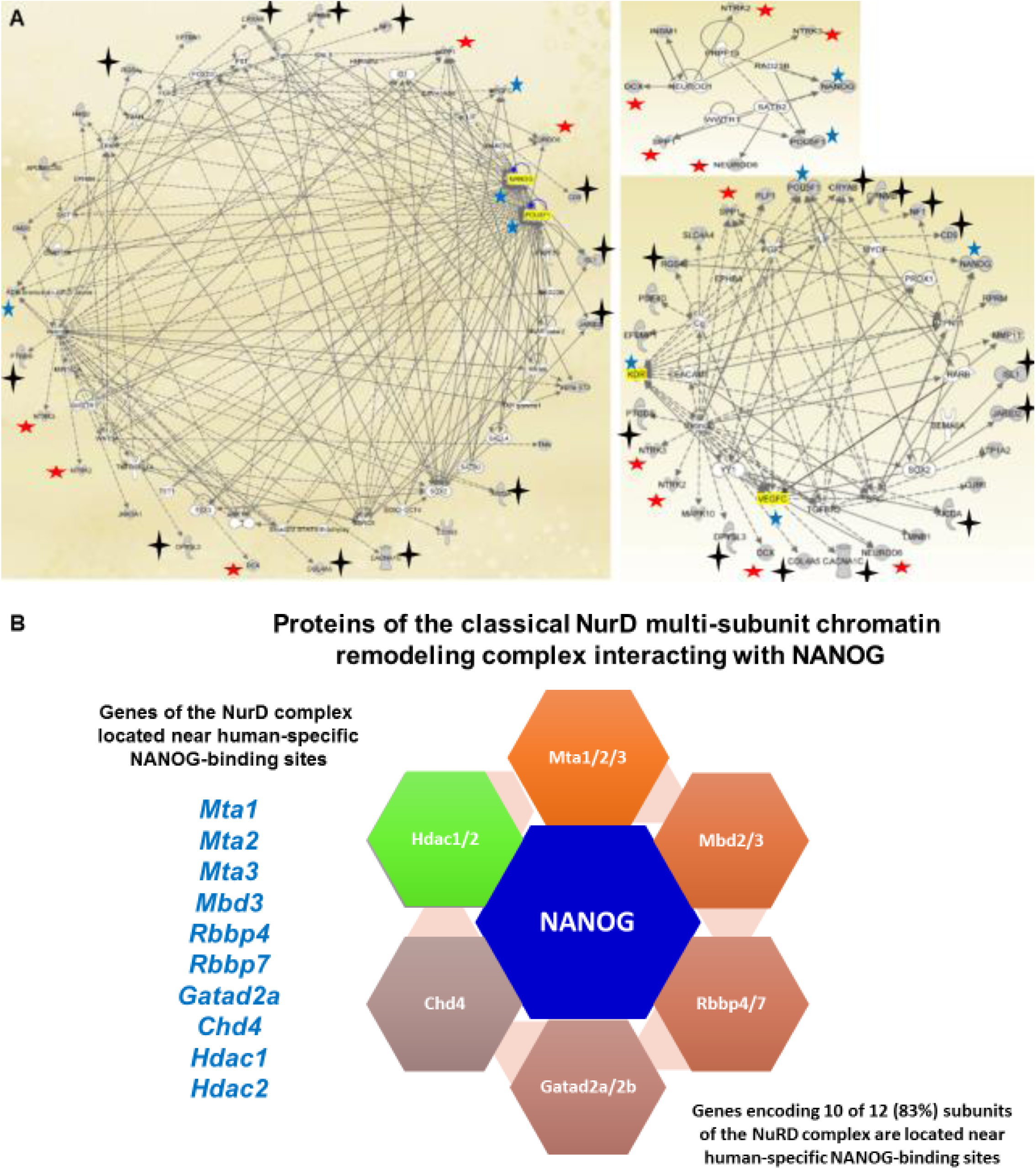
Genomic regulatory networks (GRNs) of human neocortex/prefrontal cortex development and functions that are associated with HSNBS and under potential regulatory controls of the classical multi-subunit NurD chromatin remodeling complex. A. Ingenuity Pathway Analysis of human neocortex/prefrontal cortex signature genes identifies candidate GRNs that appear under putative regulatory controls of the NANOG and POU5F1 proteins in embryogenesis and seem to transition to regulatory control of the VEGFC and its receptor KDR at the later developmental stages and in the adult brain. See text for details. B. All 12 proteins comprising the classical multi-subunit NurD chromatin remodeling complex represent protein partners of both NANOG-centered and POU5F1-centered protein-protein interaction networks in hESC (van den Berg et al., 2011; Gagliardi et al., 2013). Genes encoding 10 of 12 (83%) subunits of the classical multi-subunit NurD chromatin remodeling complex are located near human-specific NANOG-binding sites in the hESC genome. See text for additional details.

One of the intriguing features seemingly connecting these networks is that some genes identified as potential targets within one network appear within another network as the regulatory hubs, which are marked as the blue five-pointed stars in the Fig. 5. This observation suggests that these networks are interconnected by the positive feedback regulatory loops, which appear designed to support the sustained networks’ activity. The largest network is governed by the NANOG and POU5F1 proteins, which are also identified as potential targets of the second regulatory network. The *VEGFC* and *KDR* genes appear as the main regulatory hubs of the second putative regulatory network, within which the *NANOG* and *POU5F1* are depicted as potential target genes (Fig. 5). Conversely, the *VEGFC* and *KDR* genes were identified as the putative targets of the regulatory network governed by the NANOG and POU5F1 proteins (Fig. 5).

Taken together, the results of this analysis support the hypothesis that during embryogenesis the NANOG and POU5F1 proteins initiate the gene expression changes characteristic of neocortex and prefrontal cortex regions of human brain, which are discernable as a highly concordant 4,957-gene expression signature. At the later stages of brain development and functioning during infancy and transition to adulthood, the VEGFC and its receptor KDR may be responsible, in part, for maintenance and sustained activity of the fetal gene expression program in the neocortex/prefrontal cortex regions. This conclusion opens attractive opportunities for targeted pharmacological interventions using small molecule agonists and/or antagonists of the VEGFC/KDR axis *in vivo*.

### Placement enrichment analysis of HSNBS near genes encoding the protein partners of the NANOG-centered protein interaction network in embryonic stem cells

One of the key mechanisms by which NANOG delivers its critical regulatory functions in ESC is the formation of protein-protein complexes with numerous protein partners, which are designated collectively as the NANOG interactome (Wang et al, 2006; Wu et al, 2006; Liang et al, 2008; Costa et al, 2013; Gagliardi et al., 2013). Recent high-definition analysis of physically interacting proteins in ESC using an improved affinity purification protocol expanded the NANOG interactome to more than 130 proteins (Gagliardi et al., 2013), many of which are components of large multi-subunit complexes involved in chromatin remodeling. Placement enrichment analysis identifies 58 genes encoding protein-protein interaction partners of NANOG that are located in close proximity to HSNBS defined by the 1.5 Mb placement enrichment metric. Genetic components of the multi-subunit chromatin remodeling complex NuRD appear most significantly overrepresented among genes encoding NANOG-interacting proteins: genes encoding ten of twelve NuRD complex protein subunits (83%) are located in close proximity to HSNBS (Fig. 5). Notably, 27 of 58 genes encoding protein partners of NANOG that are located near HSNBS have been previously identified as the components of the POU5F1 (OCT4)-centered protein interaction network in ESC (van den Berg et al., 2011). Significantly, the efficient association in the ESC nucleus of both POU5F1 (OCT4) and NANOG proteins with all twelve proteins comprising known subunits of the classical NuRD complex has been documented (van den Berg et al., 2011; Gagliardi et al., 2013), which indicates that association of both NANOG and POU5F1 proteins with classical NuRD complex may constitute an important mechanism of their biological activity in ESC.

Collectively, the result of these analyses strongly argue that genes encoding proteins that physically interact with both NANOG and POU5F1 in the ESC nucleus may represent functionally-relevant targets of the HSGRL-associated regulatory networks. Genetic components of the classical multi-subunit chromatin remodeling complex NuRD seem particularly relevant in this regard and they should constitute the primary candidates for the follow-up structural-functional analyses and mechanistic experiments aiming to dissect their roles in human brain development and functions. Consistent with this hypothesis, recent experiments directly implicated the NuRD chromatin remodeling complex in establishing synaptic connectivity in the rodent brain (Yamada et al., 2014) and in directing the timely and stable peripheral nerve myelination by Schwann cells in mice (Hung et al., 2012). Furthermore, significant age-associated decline of expression in dentate gyrus sub-region of the hippocampus of one of the NuRD complex subunits, RbAp48 (RBBP4), was mechanistically connected with the age-related memory loss in humans (Pavlopoulos et al., 2013).

## Discussion

It has been hypothesized that one of the principal biologically-significant functions of HSGRL is the creation of new enhancer elements, leading to the increased density of conventional enhancers and facilitating transition to SED structures (Glinsky, 2016; 2017). Potential mechanisms of HSGRL-mediated effects on principal regulatory structures of interphase chromatin are likely involve emergence of overlapping CTCF/cohesin sites and LMNB1-binding sites as well as continuing insertion of Alu clusters near the putative DNA bending sites of the chromatin looping structures (Glinsky, 2017). Collectively, the ensemble of these structural changes facilitated by the targeted placements and retention of HSGRL at specific genomic locations would enable the emergence of new SED structures and remodeling of existing TADs to drive structural-functional evolution of genomic regulatory networks. These models of dynamic transitions between the interphase chromatin principal regulatory structures are in accord with the results of recent high-resolution in situ Hi-C experiments demonstrating that human genomes are partitioned into contact domains consisting of ~10,000 loops accommodating functional links between enhancers and promoters (Rao et al., 2014). Consistent with the idea that chromatin loops frequently demarcate the boundaries of contact domains, anchors at the loop bases typically occur at the contact domain boundaries and involve binding of two CTCF/cohesin sites in a convergent orientation with the asymmetric binding motifs of interacting sites aligned to face each other (Rao et al., 2014).

The potential practical utility of these concepts are illustrated by building the working models of *NANOG*, *POU5F1*, and *POU3F2* super-enhancers’ domains and associated TAD structures in the hESC genome (Figs. 1-3; Supplemental Figs. S1, S2). Combination of the utility functions of the UCSC Genome Browser facilitated visualization of key structural elements of potential functional significance within each model of putative SED structures (Figs. 1-3; Supplemental Figs. S1; S2). Utilizing structural-functional guidance derived from these models, conserved profiles and uniquely distinct patterns of H3K27ac peaks’ distribution within the *NANOG* locus-associated genomic region in 43 types of human cells and tissues were inferred (Fig. 3).

Putative regulatory consequences of targeted placements of HSGRL on expression of protein-coding genes were assessed during the analysis of expression profiles of protein-coding genes implicated in the development of the fetal and adult brain (Johnson et al., 2009; Zhang et al., 2011). These analyses indicate that HSNBS may have contributed to the establishment during the embryogenesis of the genome-wide expression changes characteristic of neocortex and prefrontal cortex regions of human brain, which are retained as a highly concordant 4,957-gene expression signature for many years of human development and maintained throughout the adulthood (Fig. 4). The follow-up Ingenuity pathway analysis confirmed that NANOG and POU5F1 proteins are most likely candidates to initiate the gene expression changes characteristic of neocortex and prefrontal cortex regions of human brain (Fig. 5), which are discernable as a highly concordant 4,957-gene expression signature. At the later stages of brain development and function during infancy and transition to adulthood, the VEGFC and its receptor KDR may be responsible, in part, for maintenance and sustained activity of the fetal gene expression program in the neocortex/prefrontal cortex regions.

Strikingly, placement enrichment analysis of HSNBS near genes encoding the protein partners of the NANOG-centered protein interaction network in embryonic stem cells identified ten genes encoding protein subunits of the classical NuRD multi-subunit chromatin remodeling complex as the candidate principal genomic regulatory elements, activity of which appears preferentially targeted by insertions of HSRGL. Importantly, all these regulatory proteins were previously identified as the protein-protein interactions partners of both NANOG and POU5F1 (OCT4) proteins in the hESC nucleus (van den Berg et al., 2011; Gagliardi et al., 2013), thus, supporting the idea that activities of NANOG and POU5F1 proteins via engagement of the classical NuRD multi-subunit chromatin remodeling complex play a central role in defining key human-specific elements of gene expression changes during the embryonic development of human brain.

## Concluding remarks

Recent analyses indicate that targeted placements and/or retention of thousands HSGRL may have a major biologically-significant effects on the principal regulatory structures of interphase chromatin in hESC, namely TADs, SEs, and SEDs (Glinsky, 2016; 2017). The key creative events in this continuing chromatin domain architecture remodeling process is the HSGRL-enabled emergence of new enhancer elements, increasing density of which would increase the probability of structural transition from conventional enhancers to novel SEs and formation of new SEDs. The successful implementation of this continuing chromatin structure remodeling process would require the presence of conserved and/or newly created overlapping CTCF/cohesin-binding sites flanking novel SEs to fulfil the essential requirement for the formation of functional SEDs. This evolutionary process involves continuing removal of old and creation of new TFBS through multiple trial-and-error events enabled by retrotransposition, cytosine methylation-associated DNA editing (MADE), and recombination (Glinsky, 2015-2017. Importantly, these HSGRL-associated changes of the nuclear regulatory architecture may facilitate the enhanced precision of regulatory interactions between enhancers and target genes within reconstructed insulated SED neighborhoods and rewired TAD structures.

Recently reported enhancers’ half-lives and mean-lifetimes (Villar et al., 2015) were estimated at 296 and 427 hundred million years, respectively. Comparisons of estimates of time periods required for creation of enhancers and SEs (Glinsky, 2017) with these estimates seem to indicate that new enhancer elements are created at a markedly faster pace during evolution compared with their decay time span. This significant dichotomy of time period requirements for new enhancers’ creation versus enhancers’ decay highlights an underlying mechanism for increasing genomic complexity as a major trend during evolution of genomic regulatory networks. A vast majority of distinct classes of regulatory elements in the hESC genome appears created on conserved DNA sequences (Glinsky, 2015-2017), suggesting that exaptation of ancestral DNA constitutes a main mechanism of creation of new regulatory sequences in the human genome. This conclusion is an agreement with recent reports describing exaptation of ancestral DNA as a mechanism of creation of human-specific enhancers active in embryonic limb (Cotney et al., 2013) and as a prevalent mechanism of recently evolved enhancers’ creation during the mammalian genome evolution (Villar et al., 2015).

Evolution of enhancers may represent an example of evolution of evolvability (He et al., 2012; Duque et al., 2014), according to which a genotypic feature evolves not due to its functional effect, but due to its effect on the ability of DNA to evolve more quickly (Wagner and Altenberg, 1996). In this context, the regulatory balance of HAT and HDAC activities at specific genomic locations as well as the performance of enzymatic systems regulating cytosine recovery/methyl-cytosine deamination cycles and DNA recombination processes may play a major role during the evolution of regulatory DNA sequences and emergence of genomic regulatory networks controlling unique to human phenotypes, including human brain development and functions. Critical experimental and clinical assessments of potential therapeutic opportunities for pharmacological interventions targeting these enzymatic systems and functions of VEGFC/KDR axis would be of immediate interest.

## Materials and Methods

### Data Sources and Analytical Protocols

Solely publicly available datasets and resources were used for this analysis as well as methodological approaches and a computational pipeline validated for discovery of primate-specific gene and human-specific regulatory loci (Tay et al., 2009; Kent, 2002; Schwartz et al., 2003; Capra et al., 2013; Marnetto et al., 2014; Glinsky, 2015). The analysis is based on the University of California Santa Cruz (UCSC) LiftOver conversion of the coordinates of human blocks to corresponding non-human genomes using chain files of pre-computed whole-genome BLASTZ alignments with a minMatch of 0.95 and other search parameters in default setting (http://genome.ucsc.edu/cgi-bin/hgLiftOver). Extraction of BLASTZ alignments by the LiftOver algorithm for a human query generates a LiftOver output “Deleted in new”, which indicates that a human sequence does not intersect with any chains in a given non-human genome. This indicates the absence of the query sequence in the subject genome and was used to infer the presence or absence of the human sequence in the non-human reference genome. Human-specific regulatory sequences were manually curated to validate their identities and genomic features using a BLAST algorithm and the latest releases of the corresponding reference genome databases for time periods between April, 2013 and June, 2015.

Genomic coordinates of 3,127 topologically-associating domains (TADs) in hESC; 6,823 hESC-enriched enhancers; 6,322 conventional and 684 super-enhancers (SEs) in hESC; 231 SEs and 197 SEDs in mESC were reported in the previously published contributions (Dixon et al., 2012; Xie et al., 2013; Hnisz et al., 2013; Whyte et al., 2013; Dowen et al., 2014). The primary inclusion criterion for selection of the human-specific genomic regulatory loci (HSGRL) analyzed in this contribution was the fact that they were identified in human cells lines and primary human tissues whose karyotype were defined as “normal”. The following four HSGRL families comprising of 10,598 individual regulatory DNA sequences were analyzed in this study: 1) Human accelerated regions (HARs; Capra et al., 2013); 2) Human-specific transcription factor-binding sites (HSTFBS; Glinsky, 2015); 3) hESC-derived fixed human-specific regulatory regions (hESC-FHSRR; Marnetto et al., 2014); 4) DNase hypersensitive sites-derived fixed human-specific regulatory regions (DHS-FHSRR; Marnetto et al., 2014). The number of HSGRL placed within a given TAD was computed for every TAD in the hESC genome and the HSGRL placement enrichment was calculated as the ratio of observed values to expected values estimated from a random distribution model at the various cut-off thresholds. Datasets of NANOG-, POU5F1-, and CTCF-binding sites and human-specific TFBS in hESCs were reported previously (Kunarso et al., 2010; Glinsky, 2015) and are publicly available. RNA-Seq datasets were retrieved from the UCSC data repository site (http://genome.ucsc.edu/; Meyer et al., 2013) for visualization and analysis of cell type-specific transcriptional activity of defined genomic regions. A genome-wide map of the human methylome at single-base resolution was reported previously (Lister et al., 2009; 2013) and is publicly available (http://neomorph.salk.edu/human methylome). The histone modification and transcription factor chromatin immunoprecipitation sequence (ChIP-Seq) datasets for visualization and analysis were obtained from the UCSC data repository site (http://genome.ucsc.edu/; Rosenbloom et al., 2013). Genomic coordinates of the RNA polymerase II (PII)-binding sites, determined by the chromatin integration analysis with paired end-tag sequencing (ChIA-PET) method, were obtained from the saturated libraries constructed for the MCF7 and K562 human cell lines (Li et al., 2012). Genome-wide maps of interactions with nuclear lamina, defining genomic coordinates of human and mouse lamin-associated domains (LADs), were obtained from previously published and publicly available sources (Guelen et al., 2008; Peric-Hupkes et al., 2010). The density of TF-binding to a given segment of chromosomes was estimated by quantifying the number of protein-specific binding events per 1-Mb and 1-kb consecutive segments of selected human chromosomes and plotting the resulting binding site density distributions for visualization. Visualization of multiple sequence alignments was performed using the WebLogo algorithm (http://weblogo.berkeley.edu/logo.cgi). Consensus TF-binding site motif logos were previously reported (Kunarso et al., 2010; Wang et al., 2012; Ernst and Kellis, 2013).

The quantitative limits of proximity during the proximity placement analyses were defined based on several metrics. One of the metrics was defined using the genomic coordinates placing HSGRL closer to putative target protein-coding or lncRNA genes than experimentally defined distances to the nearest targets of 50% of the regulatory proteins analyzed in hESCs (Guttman et al., 2011). For each gene of interest, specific HSGRL were identified and tabulated with a genomic distance between HSGRL and a putative target gene that is smaller than the mean value of distances to the nearest target genes regulated by the protein-coding TFs in hESCs. The corresponding mean values for protein-coding and lncRNA target genes were calculated based on distances to the nearest target genes for TFs in hESC reported by Guttman et al. (2011). In addition, the proximity placement metrics were defined based on co-localization within the boundaries of the same TADs and the placement enrichment pattern of HSNBS located near the 251 neocortex/prefrontal cortex-associated genes, which identified the most significant enrichment of HSNBS placement at the genomic distances less than 1.5 Mb with a sharp peak of the enrichment p value at the distance of 1.5 Mb (Supplemental Fig. S3).

The comprehensive database of expression profiles of protein-coding genes implicated in the development of the fetal and adult brain of *H. sapiens* was obtained from the previously published contribution (Johnson et al., 2009; Zhang et al., 2011). Analysis of the evolutionary age of genes comprising the 4,957-gene expression signature of the neocortex/prefrontal cortex regions of human brain was carried-out by segregating genes into thirteen sub-groups based on their respective evolutionary age, ranging from 0 (oldest genes) to 12 (youngest genes) as previously defined by Zhang et al. (2011). The gene expression enrichment factors were calculated for each individual evolutionary age sub-group by comparisons of corresponding gene age-associated distribution metrics, which were derived from the analysis of gene age-associated distribution profiles of all 19,335 genes interrogated in gene expression profiling experiments and 12,885 genes with expression changes significantly different in fetal versus adult brain. The resulting values for each evolutionary age sub-group of genes were normalized to the corresponding numerical values obtained for all genes within the corresponding set.

The assessment of conservation of HSGRL in individual genomes of 3 Neanderthals, 12 Modern Humans, and the 41,000-year old Denisovan genome (Reich et al., 2010; Meyer et al., 2012) was carried-out by direct comparisons of corresponding sequences retrieved from individual genomes and the human genome reference database (http://genome.ucsc.edu/Neandertal/). Direct access to the specific Genome Browser tracks utilized for analyses and visualization: http://genome.ucsc.edu/cgi-bin/hgTracks?db=hg18&position=chr10%3A69713986-69714099&hgsid=393865029yg7UixUE4a4awjjTahns4KTPkIl1.

Recombination rates were downloaded from the HapMap Project (The International Hapmap Consortium, 2007) and the numbers of DNA segments with the recombination rates of 10 cM/Mb or greater were counted. This threshold exceeds ~10-fold the mean intensity of recombination rates in telomeric regions, which were identified as the regions with the higher recombination rates in the human genome. It is well known that over large genomic scales, recombination rates tend to be higher in telomeric as compared to centromeric chromosomal regions. In telomeric regions, the mean detected hotspot spacing is 90 kb and the mean intensity (total rate across the hotspot) per hotspot is 0.115 cM, whereas for centromeric regions the mean spacing is 123 kb and the mean intensity is 0.070 cM (The International Hapmap Consortium, 2007).

### Statistical Analyses of the Publicly Available Datasets

All statistical analyses of the publicly available genomic datasets, including error rate estimates, background and technical noise measurements and filtering, feature peak calling, feature selection, assignments of genomic coordinates to the corresponding builds of the reference human genome, and data visualization, were performed exactly as reported in the original publications and associated references linked to the corresponding data visualization tracks (http://genome.ucsc.edu/). Any modifications or new elements of statistical analyses are described in the corresponding sections of the Results. Statistical significance of the Pearson correlation coefficients was determined using GraphPad Prism version 6.00 software. The significance of the differences in the numbers of events between the groups was calculated using two-sided Fisher’s exact and Chi-square test, and the significance of the overlap between the events was determined using the hypergeometric distribution test (Tavazoie et al., 1999).

## Supplemental Information

Supplemental information includes Supplemental Figures S1–S5 and can be found with this article online.

## Author Contributions

This is a single author contribution. All elements of this work, including the conception of ideas, formulation, and development of concepts, execution of experiments, analysis of data, and writing of the paper, were performed by the author.

## Acknowledgements

This work was made possible by the open public access policies of major grant funding agencies and international genomic databases and the willingness of many investigators worldwide to share their primary research data. I would like to thank my colleagues for their valuable critical contributions during the informal review and formal peer review process of this work.

